# Host-parasite interactions of an avian blood parasite elucidated by single-cell transcriptomics

**DOI:** 10.1101/2025.03.14.643032

**Authors:** Tim Maximilian Rapp, Tobias Lautwein, Robert Peuß, Joachim Kurtz, Oliver Krüger, Nayden Chakarov

## Abstract

Parasites and hosts perform fine-tuned cellular and molecular interactions, known from few well-explored model systems. Recent methodological advances now allow such deep insights into non-model parasites, their hosts, and their intense reciprocal manipulation. We applied single cell RNA-sequencing to blood samples of a long-lived bird host, infected at high and low intensity with malaria-related parasites. We identify diverse parasitic traits for immune evasion and potential manipulation of the avian host immune response. The transcriptomic response of immune cells was decoupled from the infection load, consistent with evidently low pathogenicity. Changes of immune cell populations showed low activation of adaptive immunity, such as clonal expansion of lymphoid cells. This absence of antagonism indicates immunosuppression by the parasite, which may mitigate virulence and thereby facilitate host-parasite coadaptation. The identified cell-type specific markers provide an important reference and will help to understand the evolutionary forces that shape host immunity in birds. Our study illuminates in rare molecular detail a benign host-parasite interaction and bridges a large coevolutionary knowledge gap of the gradient from parasitism to commensalism and mutualism.

## Introduction

Parasites represent a significant proportion of biodiversity and have evolved to manipulate host defences in diverse ways^1–3^. From an ecological perspective, this variability of fitness effects ranges from mutualism to parasitism and is maintained by trade-offs between parasite reproduction, transmission and damage to the host (i.e. virulence)^4–9^. Conversely, host immune responses cause costs to parasites which vary strongly among and within species^10^. For example, possible infection courses of human malaria range from clearance, over asymptomatic and uncomplicated forms, to severe malaria and death^11^.

The complex lifecycles of such intracellular protozoans involve the invasion of host cells and evasion of the immune response^12–15^. In well-studied malaria parasites (genus *Plasmodium*), these include antigenic variation, dysregulation of B cells, anergy of T-cells and resulting immunosuppression^13,14^. In comparison, malaria and malaria-like parasites of birds are known to have high prevalence in host populations, great potential for detrimental effects and overall ecological importance, but the cellular aspects of their biology are largely unexplored^16^. Here, we focus on *Leucocytozoon*, distinct relatives of the blood parasite sister genera *Plasmodium* and *Haemoproteus*^17^. High host specificity of *Leucocytozoon* results in more than hundred described morphospecies and many more evolutionary lineages, each specialised on a small range of bird hosts^18^. In birds of prey, a number of specialised but disparate *Leucocytozoon* lineages indicate a long history of coadaptation^19^. Here, the parasites achieve high prevalence and infection intensity, while displaying apparent low virulence^20–23^. In an intensively studied population of raptors in Germany, *Leucocytozoon* occur at a prevalence of up to 86% in nestlings (unpublished data). Together with previously described fitness effects in birds of prey, this system allows the study of coadaptation down to the cellular and molecular levels.

Unlike other haemosporidians, *Leucocytozoon* infect diverse hosts cell types, including different leukocytes^18^. The parasitic life cycle includes merozoite stages which invade erythroblasts, erythrocytes and mononuclear leukocytes within hosts. Macro- and microgametocytes (“female” and “male”-type precursors, respectively) then develop in the infected cells. Gametogenesis is initiated upon ingestion by dipteran vectors, specifically blackflies (Simuliidae)^18^. The remarkable morphological and physiological changes of these parasites and the host cells they infect have so far never been linked to context-dependent transcriptional changes.

Very few studies have investigated the transcriptomic responses to blood parasite infections in wild birds at all and those that have only analysed whole blood transcriptomes so far, potentially convoluting cell-type specific expression changes and interactions^24–27^. Meanwhile, single cell RNA-sequencing (scRNA-seq) has emerged as one method to fill this knowledge gap, with first advances in mapping immune cells in two domestic bird species^28–30^. We extend this approach to a wild host-parasite system to identify intra- and intercellular interactions between peripheral blood cells of the common buzzard (*Buteo buteo*) and its intracellular avian malaria-like blood parasite *Leucocytozoon toddi*. By examining the effects caused by these parasites, we highlight potential mechanisms leading to immunosuppression, thereby increasing host tolerance and hence suppressing fitness costs.

## Materials & Methods

### Sample acquisition and preparation

Samples were collected in June 2021 in a population of Common Buzzards (*Buteo buteo*) in eastern Westphalia, Germany (8°25′E and 52°06′N). Occupied nests had previously been located in March and April. In a first sampling event, blood samples were taken from the ulnar vein of nestlings. Air-dried blood smears were prepared in the field, fixed in absolute ethanol and stained with Giemsa^18^. Parasitaemia of *Leucocytozoon* was estimated microscopically based on gametocyte counts. From this pool of 345 buzzard nestlings sampled from 164 territories, six individuals were chosen for further sampling based on their infection status (“low” versus “high” parasitaemia; Tab. S1). Uninfected nestlings were not available due to the extremely high prevalence of the parasite in the study population. The selected individuals originated from six different breeding territories, which were evenly distributed throughout the study area. In а second sampling event (maximum ten days after first sampling), three individuals with “low parasitaemia” (LP: 0.017%, 0.025% & 0.133% of total blood cells) and three individuals with “high parasitaemia” (HP: 0.956%, 1.384% & 1.699% of total blood cells) were brought to the Genomics & Transcriptomics Laboratory of the Heinrich-Heine-University in Düsseldorf, Germany. A blood sample with a volume of 1 mL was taken from the ulnar vein, transferred to a 15 mL centrifuge tube for Percoll (Sigma-Aldrich) density gradient centrifugation with discontinuous layers of 55%, 45%, 35%, and 25% solutions of stock isotonic Percoll (SIP) diluted with PBS at 7.4 pH. The density gradient centrifugation was performed to deplete erythrocytes (swinging-bucket centrifuge rotor, 500 x g, 15min at 20°C). The easily visible upper non-erythrocytic phases, containing larger and less dense cells compared to avian erythrocytes, were extracted, and immediately subjected to single cell processing. Variation in the extraction of leukocytes may cause differences in cell-type proportions between samples as highly infected samples have less discrete layering of cell-types during density gradient centrifugation. For comparison of cell-type proportions with scRNA-seq all peripheral blood cell-types were counted via microscopy of blood smears. Per sample approximately 30000 cells were counted within 20 random fields of view (Tab. S2). The nestlings were returned to their nests immediately after sampling and transport. Sampling and transport of nestlings were permitted by the ethics commission of the Animal Care and Use Committee of the German North Rhine-Westphalia State Office for Nature, Environment and Consumer Protection (Landesamt für Natur, Umwelt und Verbraucherschutz Nordrhein-Westfalen) under the reference number 84-02.04.2017.A147.

### Single Cell Library Generation

A total of ∼40,000 single cells were generated on the 10X Chromium Controller system for single-cell droplet library generation, utilizing the Chromium Single Cell 3’ NextGEM Reagent Kit v3.1 (10X Genomics, Pleasanton, CA, USA) according to manufacturer’s instructions. Sequencing was carried out on a NextSeq550 system (Illumina Inc. San Diego, USA) with a mean sequencing depth of ∼50,000 reads/cell.

### Processing of 10X Genomics single cell data

Raw sequencing data were processed using the 10X Genomics CellRanger software (v7.0). Raw BCL-files were demultipexed and processed to Fastq-files using the CellRanger *mkfastq* pipeline. A custom reference was created by combining draft *B. buteo* and *Leucocytozoon toddi* genomes using the cellranger *mkref* pipeline. Alignment of reads to the genomes and UMI (unique molecular identifier) counting was performed via the CellRanger *count* pipeline to generate a gene-barcode matrix. All samples were aggregated and normalised for sequencing depth using the cellranger *aggr* pipeline.

Further analyses were carried out with the Seurat v4.1.1 R package^31‒33^. Initial quality control consisted of removal of cells with fewer than 200 detected genes as well as removal of genes expressed in less than three cells. Furthermore, cells with a mapping rate of > 10% to the mitochondrial genome were removed, as they most likely represent dead or damaged cells. Cell doublets were removed from the dataset using DoubletFinder v2.0^34^. Normalization was carried out utilizing SCTransform. Dimensional reduction of the data set was achieved by Principal Component analysis (PCA), based on identified variable genes and subsequent UMAP (Uniform Manifold Approximation and Projection) embedding. The number of meaningful Principal Components (PC) was selected by ranking them according to the percentage of variance explained by each PC, plotting them in an “Elbow Plot” and manually determining the number of PCs that represent the majority of variance in the data set. Cells were clustered using the graph-based clustering approach implemented in Seurat. Marker genes differentially upregulated in each cluster in comparison to all other clusters (MGs; Tab. S3) as well as differentially expressed genes (DEGs) comparing LP and HP nestlings (Tab. S4) were calculated using a Wilcoxon Rank Sum test implemented in Seurat. Additionally, correlations of gene expression with parasitaemia determined by blood smear cell counts (see methods above) were performed. Linear models were fitted to the averaged expression of each gene per cell cluster versus the gradient of parasitaemia values among samples to identify parasitaemia-correlated genes (PCGs; Tab. S5). The false discovery rate was calculated to account for multiple testing.

### Cluster annotation and downstream analyses

Literature-curated markers were used to annotate Seurat clusters with avian peripheral blood cell-types and haemosporidian gametocyte stages (Fig. 1, S1 & S2). Blood cell markers from *Gallus gallus* and *Plasmodium falciparum* (lineage 3D7) gametocyte markers were utilised^30,35‒40^. The annotation of cell-types was further aided by the overall similarity of transcriptional profiles (Fig. 1A), marker genes with indicative functions (MGs; Tab. S3) and parasite transcript abundance (Fig. S3D & S4). Multiple clusters could be grouped into one cell type based on the expression of marker genes, while the identity of subtypes remained elusive. Downstream analyses were performed using R version 4.1.2 for data processing and exploration^41^.

**Figure 1:**
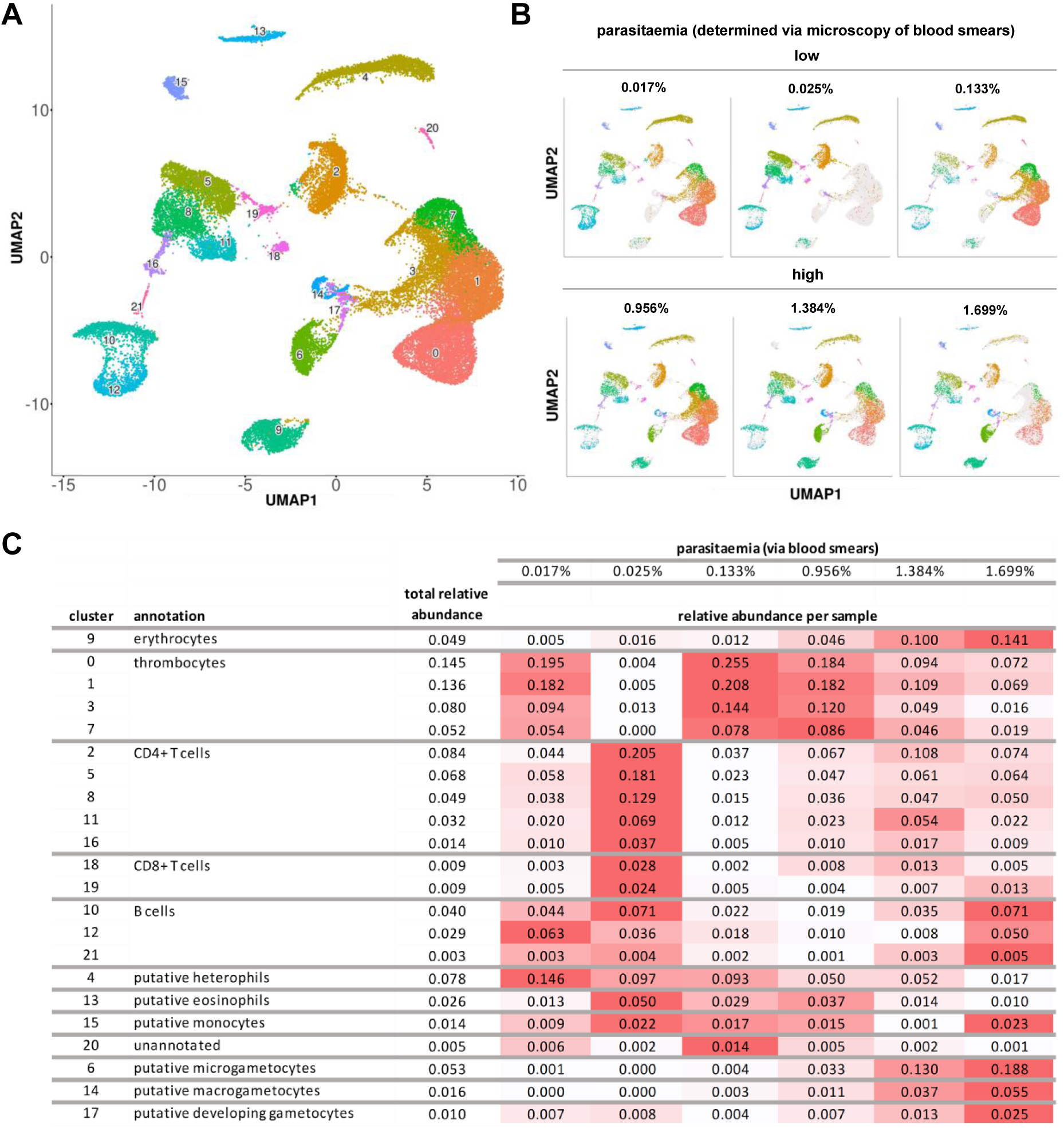
Overview of scRNA-seq clusters (Seurat clusters) and their proportions. (A) UMAP of 22 identified clusters. (B) Comparison of 6 samples categorised into LP versus HP based on their parasitaemia quantified via microscopy of blood smears. (C) Relative abundance of cell-types comparing samples ordered by parasitaemia as quantified via microscopy of blood smears. Abundance is colour-coded by a red colour scale (normalisation per row).

### Network analysis of DEGs

In the course of our analyses, we identified two clusters of interest exhibiting opposite expression patterns of the same host DEGs. We therefore performed protein-protein-interaction (PPI) network analysis of these genes. with the Cytoscape stringApp v2 using *Aquila chrysaetos chrysaetos* as reference species. Relevant DEGs for PPI networks were subsetted based on the cut-off values p ≥ 0.05 (Bonferroni adjusted) and log_2_FC (averaged per cluster) ≥1^42–44^. We chose this threshold to highlight the host’s pathways most strongly responding to changes in parasite loads. Only interactions with a confidence score of ≥ 0.9 were retrieved. Pathway enrichment analysis was performed via the enrichment tool implemented in stringApp v2 using the referenced genome as a background. For comparability, only enriched KEGG (Kyoto Encyclopedia of Genes and Genomes) pathways were highlighted.

## Results

### Identification of 22 distinct cell populations in *L. toddi*-infected peripheral blood

The transcriptomes of 38295 cells from six buzzard individuals were separately sequenced and analysed, resulting in the identification of 22 cell populations (clusters) via dimensionality reduction of transcriptional profiles (Fig. 1A). A comparison of three HP (high parasitaemia) with three LP (low parasitaemia) samples allowed the identification of the transcriptional profiles of infected cells (Fig. 1B). Literature-curated marker genes (Fig. S1 & S2) and marker genes identified by their differential expression in each cluster compared to all other clusters (MGs; Tab. S3) were used to identify the blood cell type of a given cluster. The transcriptomic similarity of parasite-infected cells and thrombocytes was high, causing these cell types to cluster together. Thrombocytes might hence be the most common host cell type of *Leucocytozoon* in buzzards (Fig. 1A, mitochondrial cytochrome b *Leucocytozoon* lineage MILANS04). Overall, 28337 annotated and 712 unannotated genes were detected. Transcript diversity (Fig. S3A), expression levels (normalised UMI counts; Fig. S3B), and proportion of mitochondrial and parasite transcripts varied clearly between cell clusters (Fig. S3C-D & S4).

### Immune cell populations between infection categories

We compared LP and HP and found an expected strong increase in gametocyte proportions. This was accompanied by slightly decreasing proportions of most other cell types (Fig. 1). Signs of adaptive immunity activation, such as clonal expansion of lymphoid cells, were not found. Strong decreases were observed in heterophils and B cells in high infection intensity samples.

### Strong differential gene expression occurs in two parasite cell types

We compared gene expression between LP and HP groups to identify expression patterns responding to differences in parasite load. Between infection categories, 2033 differentially expressed genes (DEGs) were found −1138 parasite DEGs, 866 host DEGs and 29 unannotated DEGs (Tab. S4, Fig. 2 & S2; cut-off value: p-value (adjusted) ≤ 0.05). We subsetted the strongest DEGs to highlight the most influential changes between low and high infection loads (Tab. S4, Fig. 2, S5 & S6; cut-off values: p-value (adjusted) ≤ 0.05; log₂-FC (averaged) ≥ 1). In the following, gametocyte-infected cells will be referred to as developing gametocytes, micro- and macrogametocytes for simplicity. The highest number of DEGs was found in microgametocytes and developing gametocytes, while macrogametocytes and host cells generally expressed fewer DEGs (Fig. S6A-C). Among gametocyte clusters, we observed more shared DEGs between microgametocytes and developing gametocytes, than with macrogametocytes (Fig. 2A). Host DEGs were largely expressed under LP, whereas parasite DEGs were predominantly present in HP samples (Fig. 2B, S6B & S6C). This implies an overall downregulation of host transcriptomic responses under HP. Since parasite genes dominated under HP in developing gametocytes and the unannotated genes followed the same trend, they likely represent parasitic transcripts. The strongest parasite DEGs were exclusive to HP, but present in all host clusters.

**Figure 2:**
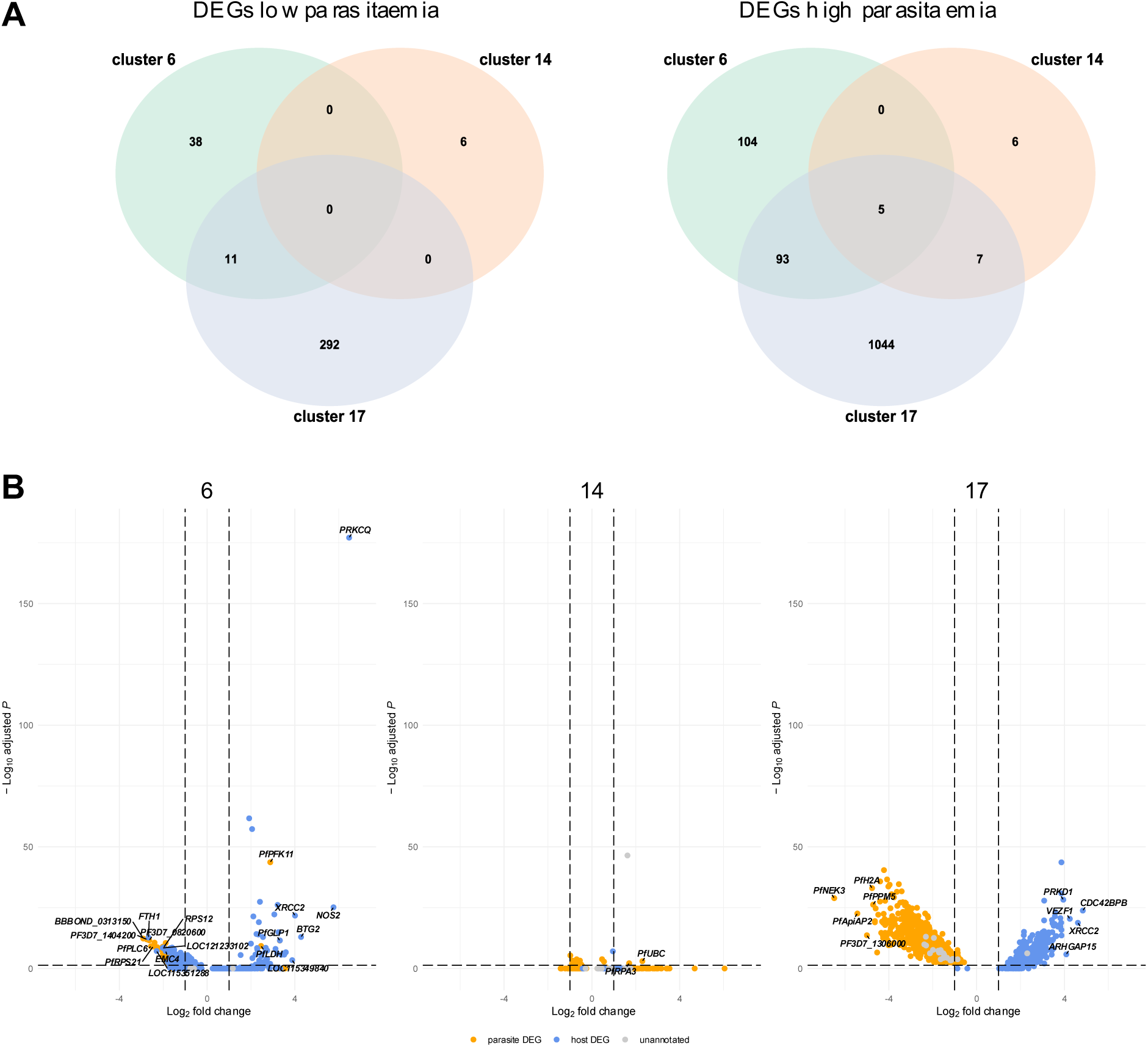
(A) Venn diagrams of DEGs in gametocyte clusters comparing LP and HP (cut-off: p-value (adjusted) ≤ 0.05). (B) Differential gene expression across gametocyte clusters (top five host and parasite DEGs of LP versus HP are labelled). Dotted lines delimit cut-off-values (p-value (adjusted) ≤ 0.05; log₂-FC (averaged) ≥ 1). Negative and positive values for log₂-FC represent differential expression under HP versus LP respectively.

A comparison of uninfected versus infected samples was not feasible due to the high prevalence of *Leucocytozoon* in the wild study population during sample acquisition. In order to validate that differential expression analysis of HP versus LP groups represented changes in gene expression related to gradual changes in parasitaemia, we correlated gene expression with parasitaemia (infection intensity) as a continuous variable determined from blood smears. The analysis delivered fewer genes of interest with a higher ratio of parasite to host genes, likely due to statistical constraints (FDR-correction). 216 PCGs (parasitaemia-correlated genes) showed a strong correlation with parasitaemia (Tab. S5, cut-off value: FDR ≤ 0.05). Overall, 182 parasite and 34 host PCGs were correlated with the gradient of infection intensity. The correlations were almost exclusively positive apart from four host genes displaying a negative correlation with parasitaemia in heterophils. Out of the 216 PCGs (Tab. S5), 19 parasite genes also represented DEGs retrieved by differential expression analysis between HP and LP (Tab. S5; cut-off value: p-value (adjusted) ≤ 0.05).

### Cell type specificity of DEGs

We assessed how specific DEGs were to the different cell types by calculating the number of clusters in which each DEG was expressed. We observed a high cell-type specificity of host DEGs, likely due to the functional differentiation of blood cell types (Tab. 1, Fig. S7). Parasite DEGs were highly specific to gametocyte subtypes, aligning with sexual maturation and differentiation (Tab. 1, Fig. S7). A subset of parasite DEGs was redundantly expressed in all peripheral blood cell types and developing gametocytes, but not in macro- and microgametocytes (Tab. 1, Fig. S7). This set of genes is most likely required for the initial widescale invasion of various host cell types by merozoites.

**Table 1:**
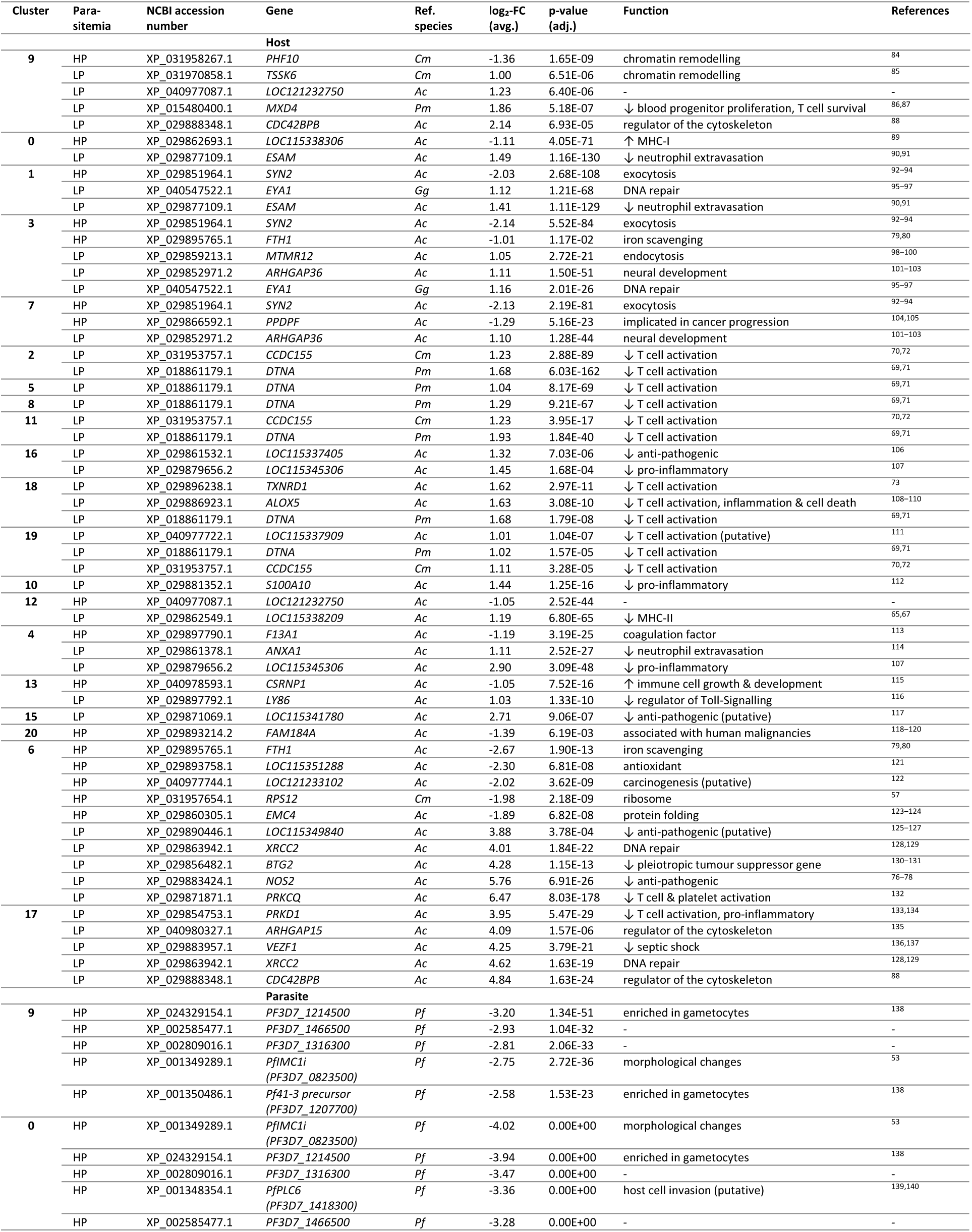
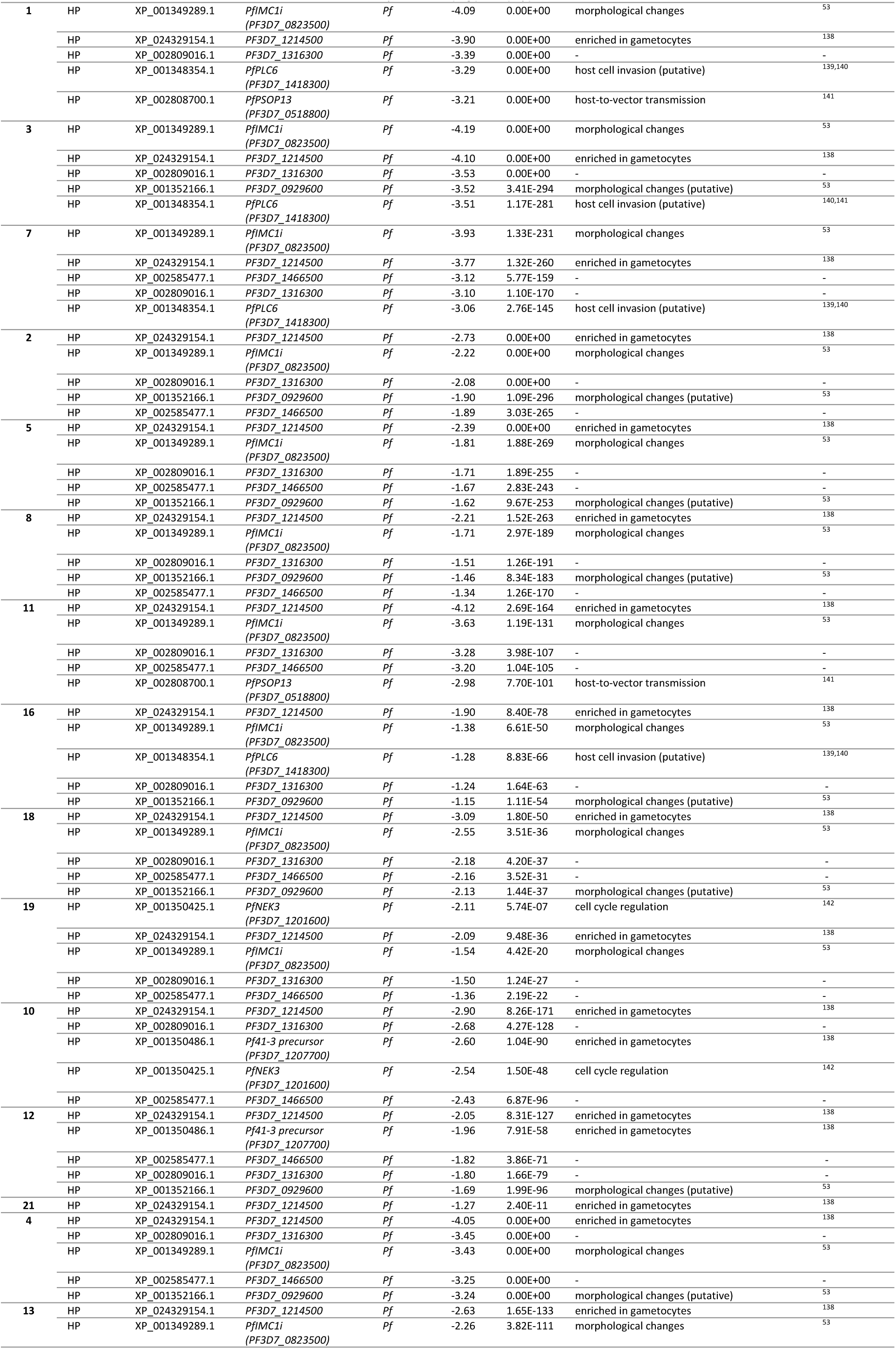
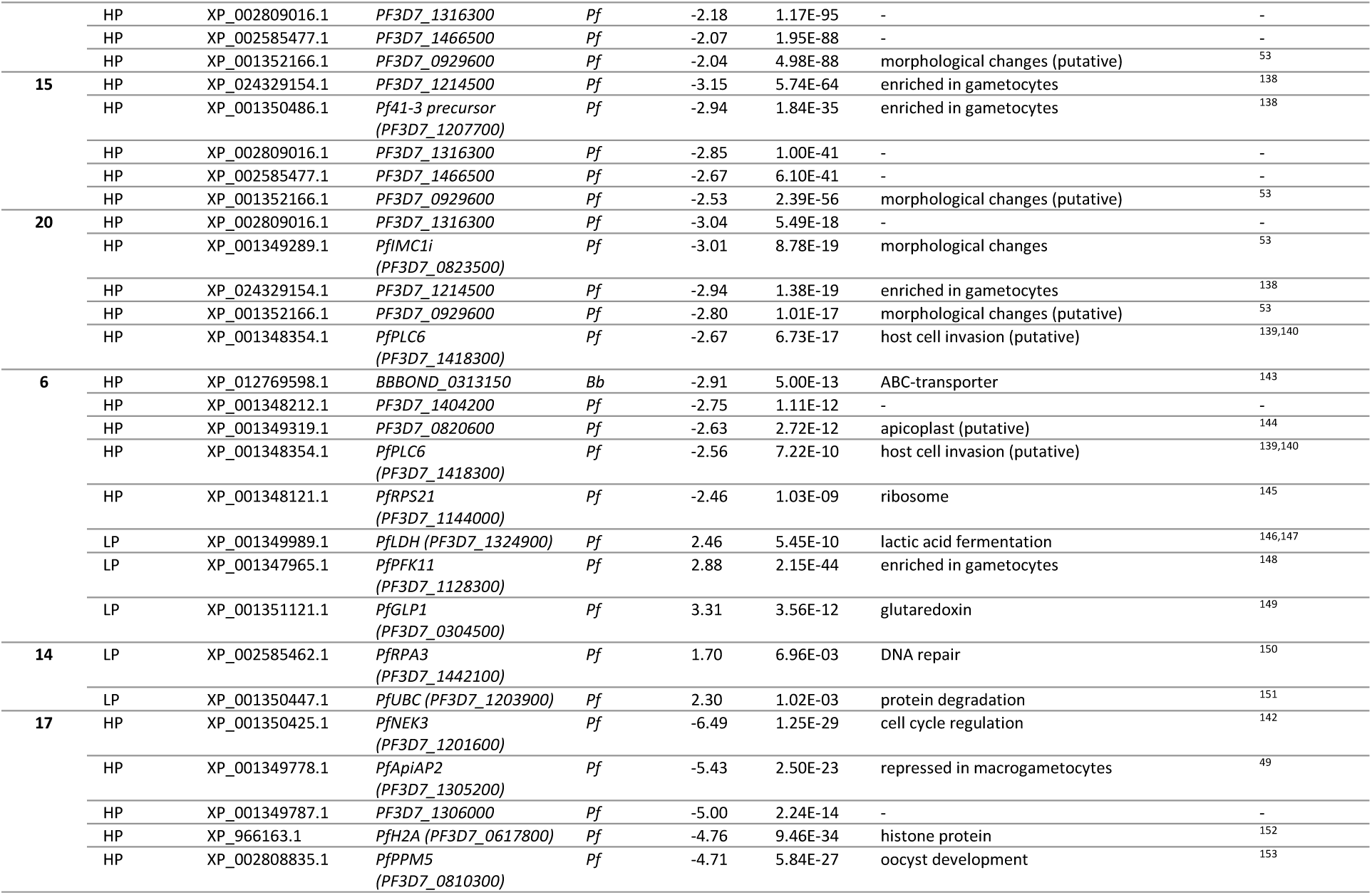
Strongest DEGs across clusters (ordered by cell type). References indicate potential gene functions. Immune genes upregulated under HP are indicated by “↑”, whereas immunosuppression (downregulation under HP) is indicated by “↓” (see column “Function”).

### Immune gene expression generally lacks correlation with infection intensity

The host immune genes which ranked among the top five DEGs in each cluster were largely downregulated under HP within various cell types (Tab. 1). These genes are implicated in processes such as in T cell activation, the inflammatory response or pathogen defence (Tab. 1).

Immune effectors, including cytokines which are crucial for the vertebrate immune response, changed little with increased parasitaemia or even displayed downregulation across cell types^45^. While generally promoting parasite elimination, an exaggerated pro-inflammatory response via TNF-α, IL-6, or IL-8 can also favour severe malaria and pathogenicity^46^. Some cytokines had weak ubiquitous expression, but were highly expressed in certain cell clusters (Tab. S6), both in HP and LP (Tab. S6). As an exception, *TGF-β* displayed differential expression under LP in thrombocytes and monocytes.

MHC-I components were upregulated under HP in thrombocytes, erythrocytes, T-cells and eosinophils (Tab. S6). In contrast, these genes were downregulated under HP in some B-cells, T-cells and in monocytes (Tab. S6). Interestingly, microgametocytes displayed upregulation of *B2M* (MHC-I component) under HP, while developing gametocytes exhibited the opposite pattern (Tab. S6). MHC-II components were downregulated in some B-cell clusters and monocytes under HP (Tab. S6). Among the detectable immunoglobulin mRNAs, only *IgL* (*Immunoglobulin light chain λ*) expressed in B-cells was significantly lower under HP (Tab. S6).

### Opposing patterns of host gene expression in gametocyte subtypes

Developing gametocytes and microgametocytes showed the highest number of DEGs (Fig. 2, S6). A subset of host DEGs in these clusters were downregulated under HP in developing gametocytes, while being upregulated under HP in microgametocytes. We therefore generated protein-protein-interaction (PPI) networks based on the strongest DEGs in these clusters to identify host pathways differently responding to infection intensity in developing gametocytes versus microgametocytes (Fig. 3). A large group of interactors with opposing expression patterns were components of the ribosome which were downregulated in developing gametocytes, but upregulated in microgametocytes under HP (Fig. 3). Additionally, an enrichment of network components involved in mRNA surveillance, the spliceosome and the regulation of actin cytoskeleton was observed in developing gametocytes (Fig. 3).

**Figure 3:**
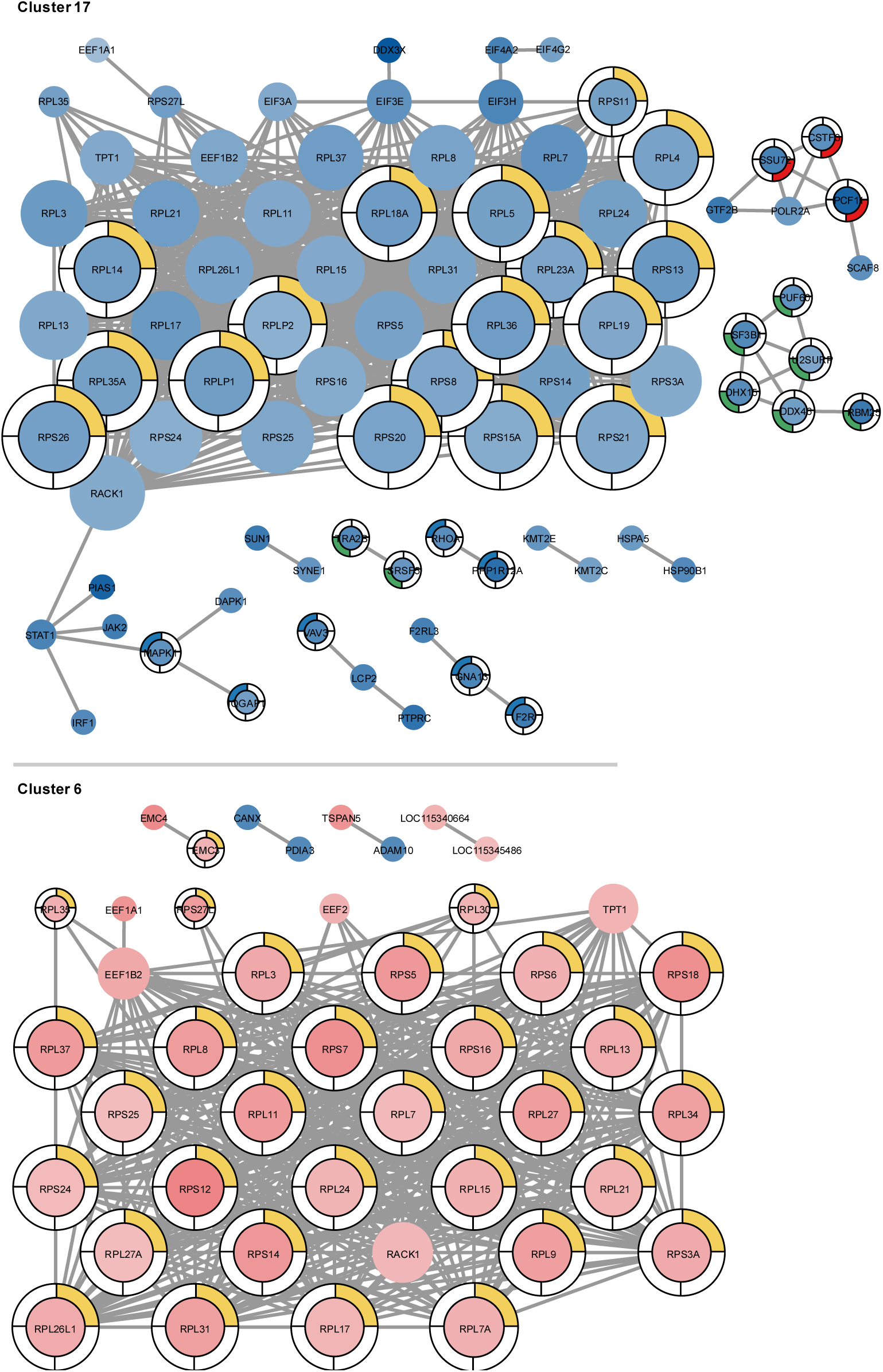
String PPI-networks of host genes (p-value (adjusted) ≤ 0.05; log₂-FC (averaged) ≥ 1) of developing gametocytes microgametocytes. Node colour intensity encodes differential expression comparing LP (blue) versus HP (red). Colouring outside nodes refers to enriched KEGG pathways (yellow: “ribosome” (False discovery rate (FDR) = 4.99E-36); red: “mRNA surveillance pathway” (FDR = 0.0451); green: “spliceosome” (FDR = 0.0047); blue: “regulation of actin cytoskeleton” (FDR = 0.0013)). Node size corresponds to number of connections.

## Discussion

This study is the first one to perform scRNA-seq on a free-living avian host. It is also the first to characterise the cells of an unculturable haemosporidian parasite in this way, demonstrating both the feasibility and opportunities of the method for parasitology, wildlife veterinary medicine and evolutionary ecology. We focused on changes of host expression linked to the intensity of infection.

The annotation of *Leucocytozoon* gametocyte clusters was based on the difference between light and intense infections and led us to subdivide these clusters into putative developing gametocytes, micro-and macrogametocytes. This division is also suggested by the bifurcation of macro- and microgametocyte clusters from a probable progenitor cell cluster, much like a pattern observed in *P. falciparum*^47^. It is striking that gametocyte subtypes displayed strong differences with regard to basic cellular properties. While general transcript abundance and the ratio of parasite versus host transcripts was highest in macrogametocytes, this cluster showed the lowest proportion of parasite mitochondrial gene transcripts. The functional differences between micro- and macrogametocytes fitting these transcriptomic differences may be linked to differing mitochondrial-to-nuclear genome ratios between *Leucocytozoon* gametocyte types^48^. They are also reminiscent of a mechanism described in *Plasmodium*, where maternally derived mRNA-pools drive zygote development^49^.

We did not find a good match to *Plasmodium* gametocyte markers. This is not surprising, given the estimated phylogenetic distance of 46-74 million years between *Leucocytozoon* and other haemosporidian genera, but this complicated cell cluster annotation^17^. Our study in comparison with diverse scRNA-seq studies on *Plasmodium* can inform about cellular markers and processes which are either common or specific to different groups of blood parasites. For instance, we observed some conserved mechanisms controlling macro- versus microgametocyte differentiation in *Plasmodium* and *Leucocytozoon*. We identified an orthologue of *DOZI*, a gene which functions as a translational repressor in macrogametocytes of *Plasmodium* and as a marker of macrogametocytes^49^. With the help of such markers, gametocyte ratios, demographic, and physiological traits of diverse haemosporidian populations may become accessible based on gross blood transcriptomic data.

Classically, gametocytes of *Leucocytozoon* were thought to develop within erythroblasts, erythrocytes, and mononuclear leukocytes^18^. Recently, it has been suggested that some *Leucocytozoon* species solely infect thrombocytes, while other species vary in their host cell choice^50‒52^. The overall similarity of transcriptional profiles between gametocytes and thrombocytes points to the latter as the preferred host cell type in *Leucocytozoon* in accipitrid hosts. Thrombocytes were clustered into four cellular subtypes. Whether these subtypes have different functions remains to be investigated. Meanwhile, parasite transcripts were expressed at low levels in several host cell types. This global expression of parasite genes across host cell types under HP might be due to a promiscuous invasion of host cells by merozoites. This interpretation is supported by the upregulation of genes required for invasion of host cells, and known from merozoites in *P. falciparum* such as *PfIMC1i* and *G2*^53^. We also observed an expression of different host cell type markers in different gametocyte subtypes. Here, microgametocytes expressed more thrombocyte markers than macrogametocytes, while the opposite pattern was observed for red blood cell markers. This may be residual expression of host genes after invasion or parasite-driven ectopic expression. If so, parasite micro- and macrogametocytes might each preferentially develop from different host peripheral blood cell types (thrombocytes versus red blood cells).

We found that host transcriptional profiles differed strongly between infection intensity levels. Cells harbouring developing gametocytes, micro- and macrogametocytes even displayed partly opposing trends of gene regulation. This indicates that developing versus differentiated gametocytes face – and in turn shape – very different regulatory environments within their host cells. The bulk of DEGs occurred in developing gametocytes, corresponding to a peak of host-parasite interactions during the onset of parasite development, when the host cell is reprogrammed to match the requirements of gametocyte development^54^. The restructuring of the host cell by intracellular protozoa is well known to elicit and require profound changes in the host’s transcriptome^55‒56^.

We detected opposing expression patterns of genes involved in host ribosome biogenesis in developing gametocytes versus microgametocytes. Deregulation of such genes has been shown to enhance cancer proliferation in model systems^57^. Genes involved in the host’s ribosome biogenesis were downregulated under HP in developing gametocytes, but were upregulated in microgametocytes, and no pattern was observed in macrogametocytes. This shows a density- or infection-course-dependent mechanism modifying host ribosome biogenesis at different parasitic stages. We also observed downregulation of host RNA metabolism, fitting to allocation or hijacking of RNA resources towards parasite gene expression. Coincidentally, also a downregulation of genes linked to the host’s actin cytoskeleton was found. This phenomenon is known from the early development of other intracellular protists and may destabilise the host cell structure allowing cellular reshaping^58‒60^.

In contrast to the strong transcriptional signs of host cell manipulation, responses indicating immune activation were weak or absent. The strong downregulation of immune genes in various cell types points towards immunosuppression under increased parasite load. The absence of clear differences in immune cell populations in response to parasitaemia supports this interpretation. Immunological notion suggests that after the invasion of a vertebrate by parasites, MHC-receptors trigger clonal expansion of T- and B-cells with specific antibodies against the pathogen. However, many intracellular parasites prevent the activation and clonal expansion of T-cells and other immune cells required for parasite clearance. *Leucocytozoon* appears to use this strategy as well^12–14^. A further point in favour of immunosuppression by *Leucocytozoon* was that a majority of differentially expressed host genes were downregulated under HP. Importantly, several effectors involved in the immune response were downregulated in multiple cell types, e.g. calgranulin- and avidin-encoding genes in CD4+ T-cells or *NOS2* in microgametocytes. These genes have been shown to be transcriptionally upregulated in response to LPS-injection in wild passerines and upon *Salmonella enterica*-infection in chicken^61‒63^. Another example is the downregulation of MHC-I and -II expression, used by *Leishmania* and *Toxoplasma*^64‒67^. A similar strategy might be the downregulation of *B2M* in developing *Leucocytozoon* gametocytes under HP, since B2M is an essential component of MHC-I. Yet, it remains puzzling that we also found the opposite effect in microgametocytes. A wide range of host cell types indeed displayed upregulation of MHC-I components under HP. However, MHC-I upregulation did not seem to evoke a strong CD8+ T-cell response, as a pronounced cytokine response was lacking and genes involved in CD8+ T-cell activation were downregulated.

The weak and promiscuous expression patterns of various cytokines indicate a mild inflammatory response, regardless of infection intensity. Additionally, this response was downregulated under HP in the case of *IL-8 like* and *TGF-β* in thrombocytes. The upregulation of *IL-8 like* under HP in eosinophils might contribute to an inflammatory response, while downregulation was observable in thrombocytes. Therefore, total *IL-8 like* expression remained stable. The expression of *IL-2, IL-16, IL-18, IL-7*, *IL-15, TNF11*, *TNF13B* and *TNF8-like* did not differ between LP and HP or correlated with infection intensity. Since we could not compare infected with uninfected samples, we cannot differentiate whether the observed expression level of these cytokines is constitutive or whether it was induced during the onset of infection without responding to increases in parasitaemia. Finally, the expression of immunoglobulins remained stable or was even downregulated under HP, in agreement with an earlier finding that antibody titres of buzzard nestlings are not linked to *Leucocytozoon* infection intensity^68^.

Within the various peripheral blood cell types, we observed highly cell-type specific transcriptional responses to parasitaemia, some of which might be reactions to manipulation by the parasite. A vast body of literature on comparable host-parasite systems indicates the functions of genes which we find to be differentially expressed in connection to infection intensity. Although these connections remain speculative, they can help to understand how and which mechanisms utilised by other intracellular protozoa for host manipulation might be used by *Leucocytozoon* and which may be so far unknown. An example is the cell type-specific immunosuppression that may be provided by the downregulation of *DTNA* and of *CCDC155* in CD4+ T-cells, potentially affecting T-cell activation. DTNA is known as a component of the DGC (dystrophin-glycoprotein complex) found in striated muscle cells^69^. CCDC155 has been characterised as a component of the LINC (linkers of the nucleoskeleton to the cytoskeleton)-complex coupling chromosomes to the cytoskeleton during meiosis^70^. Immunological synapse formation during T-cell activation involves both the LINC-complex and the DGC, parts of which were downregulated in intense *Leucocytozoon* infections^71,72^. T-cell activation might also be suppressed in CD8+ T-cells, since some genes known to be involved in this process were downregulated under HP. For instance, *TXNRD1* is upregulated during activation and clonal expansion of T-cells in response to *Leishmania major* infection^73^. Consequently, *TXNRD1*-deficiency in such models leads to increased parasitaemia^73^. Apart from differential gene expression indicating immunosuppression, we also identified some DEGs which may mediate tolerance to infection by mitigating the effects of parasite growth and load. In microgametocytes, the downregulation of *NOS2* under HP could play a role in increasing host tolerance to infection. Intracellular protists are known to downregulate iNOS as a mechanism of immune evasion as exemplified by *T. gondii* and *L. amazonensis* infections^74,75^. Inducible nitric oxide synthase (iNOS) encoded by *NOS2* catalyses the production of anti-parasitic nitric oxide (NO), which is a “double-edged sword” and at excess causes cytotoxic damage to the host^76‒78^. Therefore, downregulation of *NOS2* might be beneficial, especially under high parasite burden, and might be contributing to infection tolerance. The upregulation of *FTH1* under HP in thrombocytes could contribute to tolerance in a different way. Ferritin prevents labile iron from activating pro-apoptotic signalling pathways in hepatocytes during human malaria infections, thereby increasing host tolerance to high infection loads^79^. However, it remains questionable if iron sequestration is needed during *Leucocytozoon*-infections since they are not known to cause haemolysis and excessive release of iron as seen in *Plasmodium*-infections^18^. *FTH1* was also upregulated in microgametocytes, reminiscent of a mechanism observed in *Leishmania amazonensis*-infected macrophages, where its gene product promotes parasite growth through iron scavenging^80^.

Our findings indicate that the low virulence of *Leucocytozoon* in buzzards is associated with widescale immunosuppression in peripheral blood. At the same time, our study relied on sampling wild animals in the field and not on a strictly controlled laboratory set-up. Therefore, the sampled individuals might differ in their susceptibility to infection and represent different time points of the infection course, which may also correspond to different immunological states. Immunosuppression can prevent excessive immune responses, which can be damaging to the host (e.g. excessive cytokine production^56^). Therefore, the inhibition of such pathways during parasitic infections can be beneficial to the host as long as the overall parasite virulence is low. Indeed, costs of infection in this system have been shown to be transient and likely compensated by parental care^22^. Accordingly, infection at the nestling stage is neither predictive of strong immediate pathology, nor of later survival prospects^23^. Immune evasion by parasites can have a mitigating effect on virulence and in certain contexts assist the evolution of host tolerance^3^. Vector-mediated parent-to-offspring transmission, i.e. quasi-vertical transmission, may facilitate the evolution towards low virulence in this and similar host-parasite systems^81,82^. This pattern of transmission and parasite evolution may be a consequence of family territoriality and local confinement of parents and offspring throughout the extended parental care period^82,83^.

In this light, our study starts to fill a big knowledge gap of the molecular tools and strategies used by the wide diversity of blood parasites in wild hosts. We provide a rare example of a host-parasite-system elucidated down to the cellular and molecular mechanisms, which may contribute to reduced immunopathology and improve host-parasite coadaptation. This significantly enhances our understanding of host-parasite interactions and their diverse coevolutionary paths which still remain to be comprehensively understood in the wild.

## Acknowledgments

Sampling and processing of the samples was supported by the German Research Foundation (DFG), as part of the SFB TRR 212 (NC³) – project numbers 316099922 and 396780709 and DFG project number 398434413 to OK and NC. This work was also supported by the DFG Research Infrastructure West German Genome Center, project 407493903, as part of the Next Generation Sequencing Competence Network, project 423957469. Next Generation Sequencing and/or analyses was carried out at the West German Genome Center Düsseldorf. NC and OK received the DFG grant #DJ 154/4-4 as part of the DFG Sequencing call #2. Computational infrastructure and support were provided by the Centre for Information and Media Technology at Heinrich Heine University Düsseldorf and from the Center for Biotechnology, CeBiTec, Bielefeld University.

## Data Accessibility and Benefit-Sharing

The .fastq-files containing scRNA-seq sequences are deposited in the European Nucleotide Archive (ENA) at EMBL-EBI under accession number PRJEB73843. R-code for downstream analyses is accessible via Dryad (DOI: 10.5061/dryad.sbcc2frf2).

## Author contributions

NC and OK designed the research. NC was involved in sample acquisition. NC and TL performed laboratory procedures. TL and TMR analysed the data. TMR wrote the paper. RP and JK provided critical feedback to the analyses and the manuscript.

## Supplementary Figures

**Figure S1:**
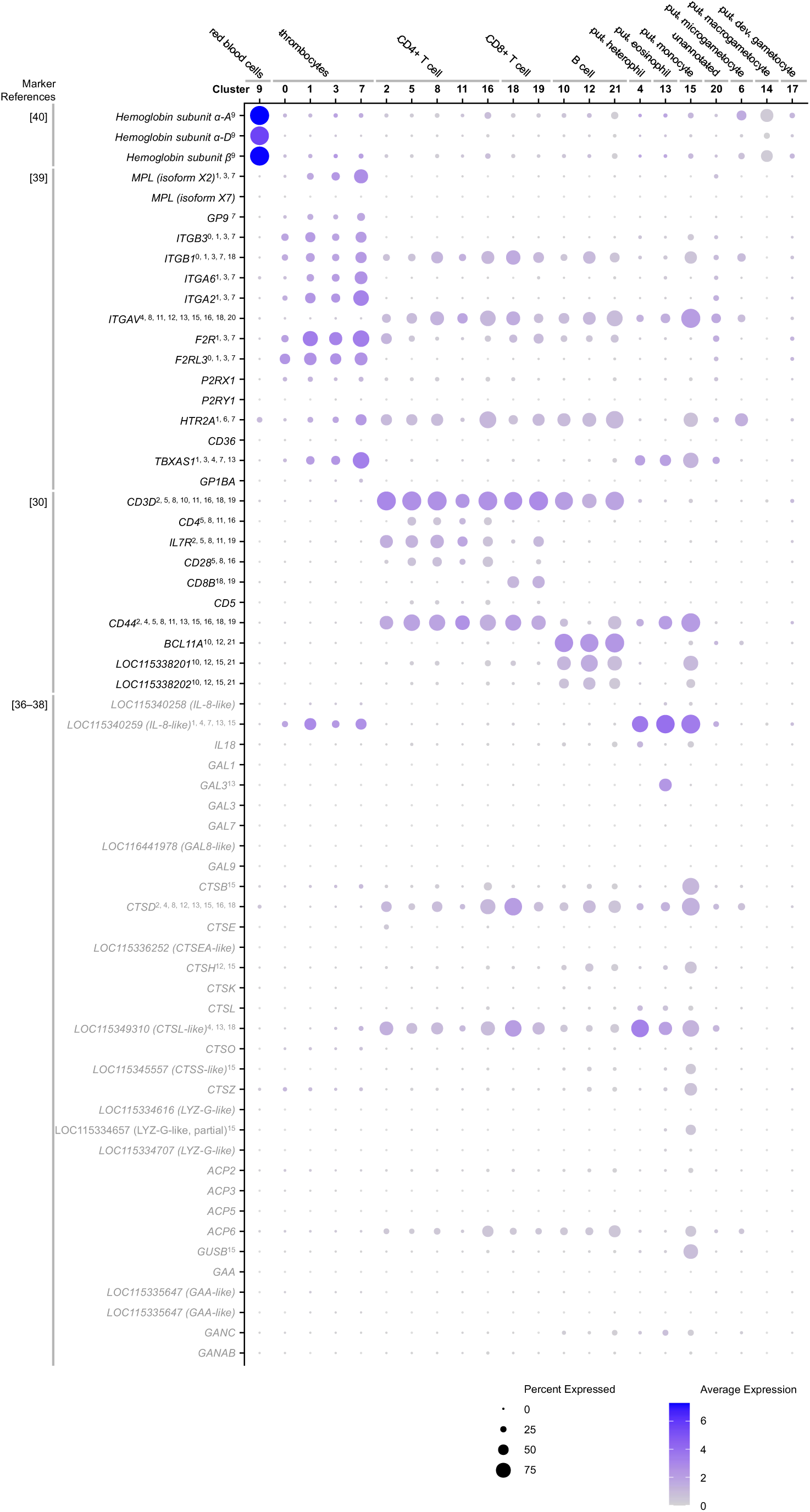
Dotplot of host marker genes curated from literature^30,36–40^. Clusters were ordered by qualitative examination of marker gene expression under consideration of cluster distribution in Fig. 1A. Final annotations are indicated at the top of the plot. Annotations in superscript list clusters in which a gene is differentially expressed in comparison to all clusters. Gene names in grey font represent indirectly inferred markers (non-transcriptomic markers). Duplicates are based on two transcript variants mapped to the same gene.

**Figure S2:**
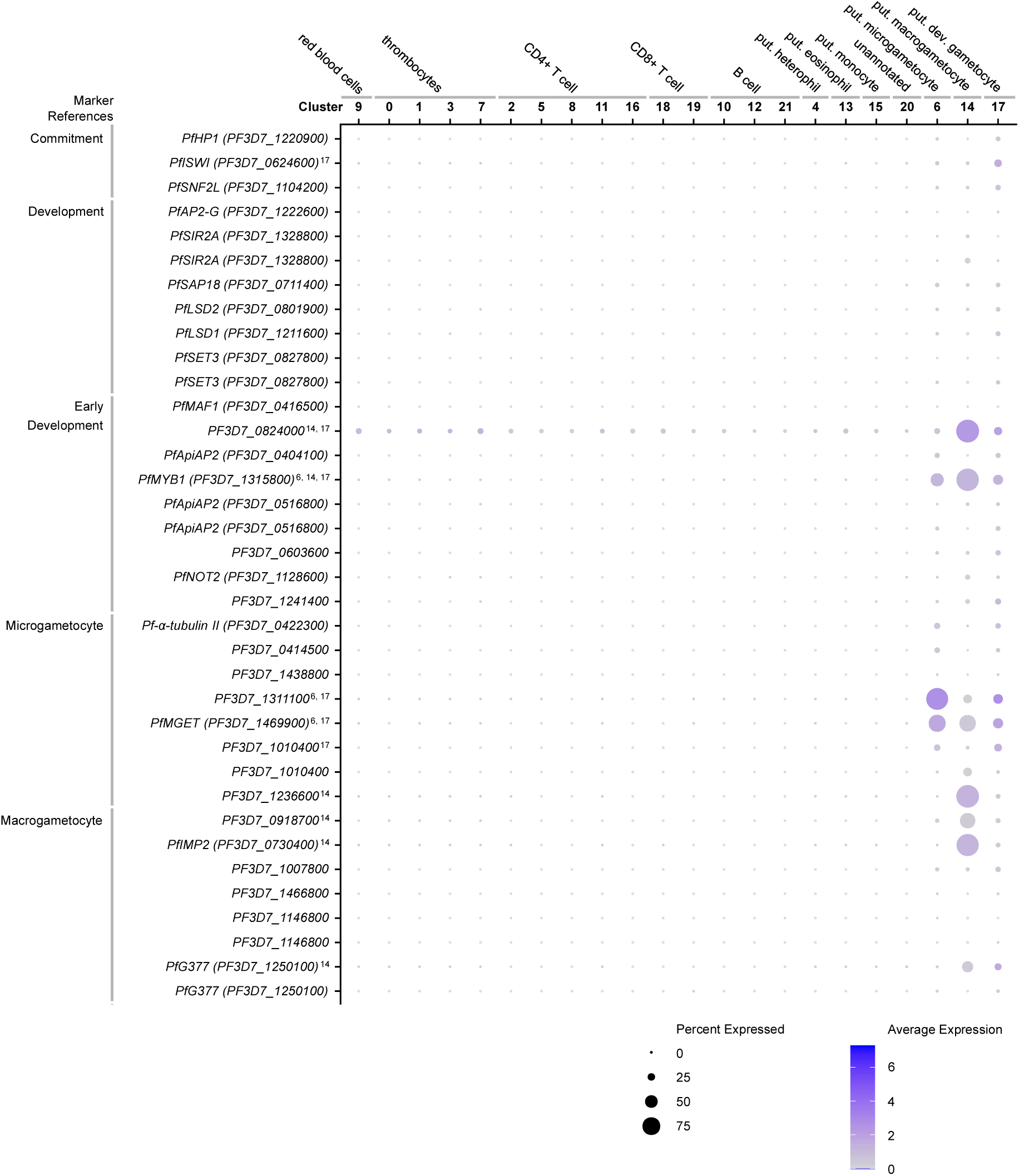
Dotplot of parasite marker genes from curated literature^35^. Clusters were ordered by qualitative examination of marker gene expression under consideration of cluster distribution in Fig. 1A. Final annotations are indicated at the top of the plot. Annotations in superscript list clusters in which a gene is differentially expressed in comparison to all clusters. Categories include commitment to gametocytogenesis (“commitment”) versus gametocytic development (“development”), early gametocyte development and micro- and macrogametocyte specific markers. Duplicates are based on two transcript variants mapped to the same gene.

**Figure S3:**
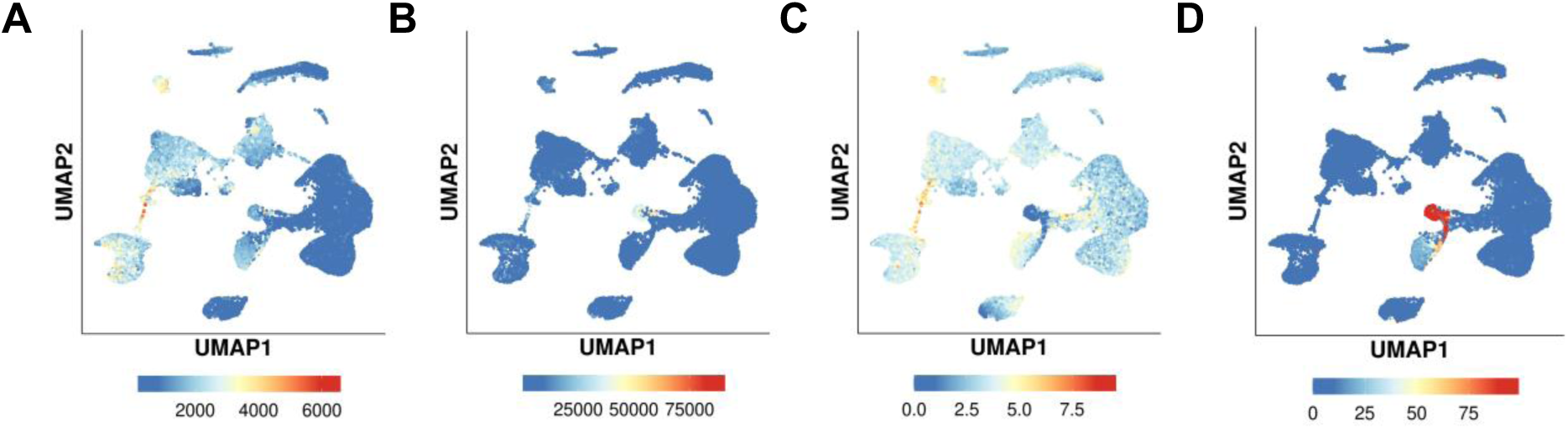
UMAPs displaying (A) number of genes expressed per cell, (B) number of UMIs detected per cell, (C) proportion of mitochondrial genes per cell and (D) proportion of parasite genes per cell (genes mapped to reference genomes of parasite species).

**Figure S4:**
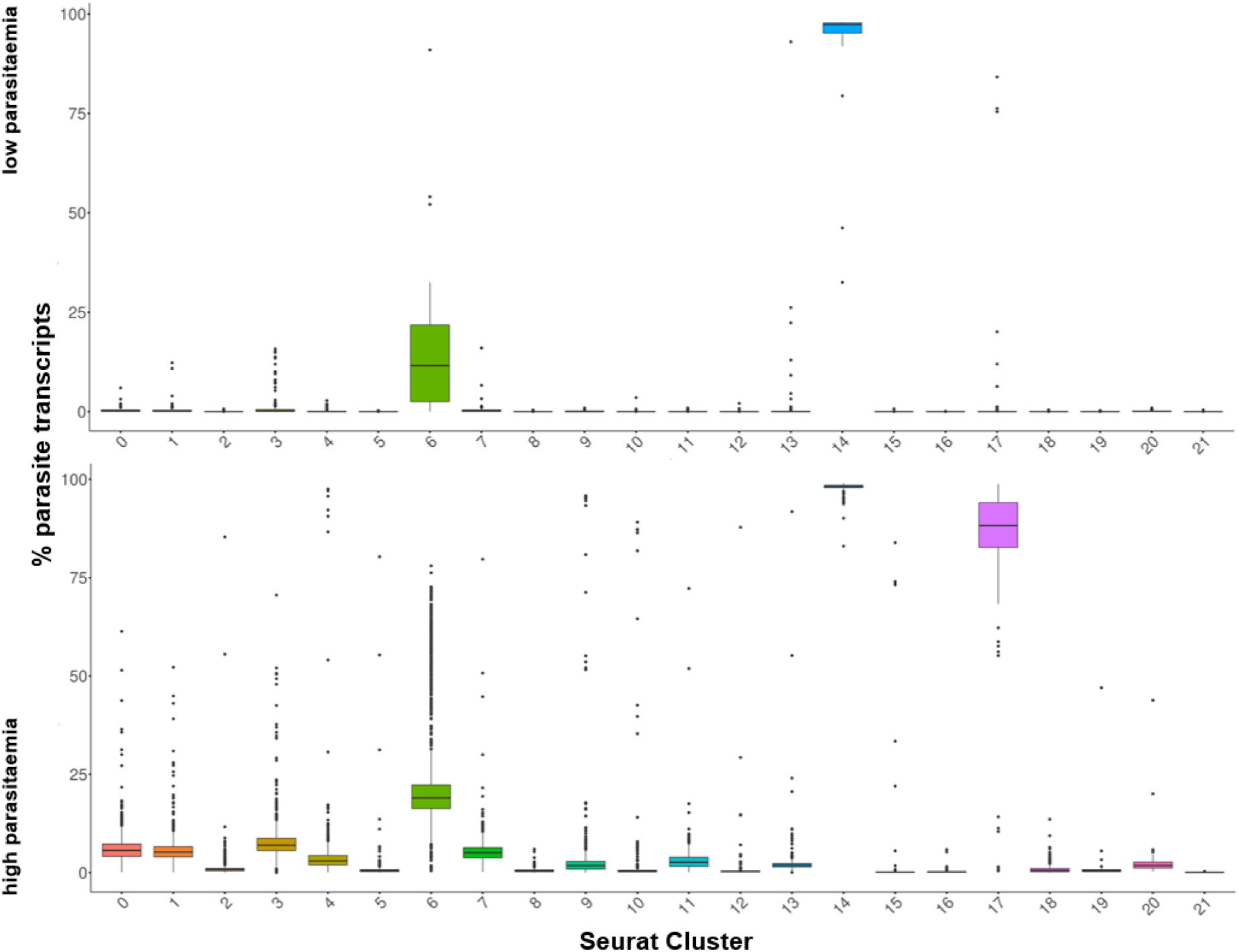
Proportion of parasite transcripts per cluster comparing low versus high parasitaemia.

**Figure S5:**
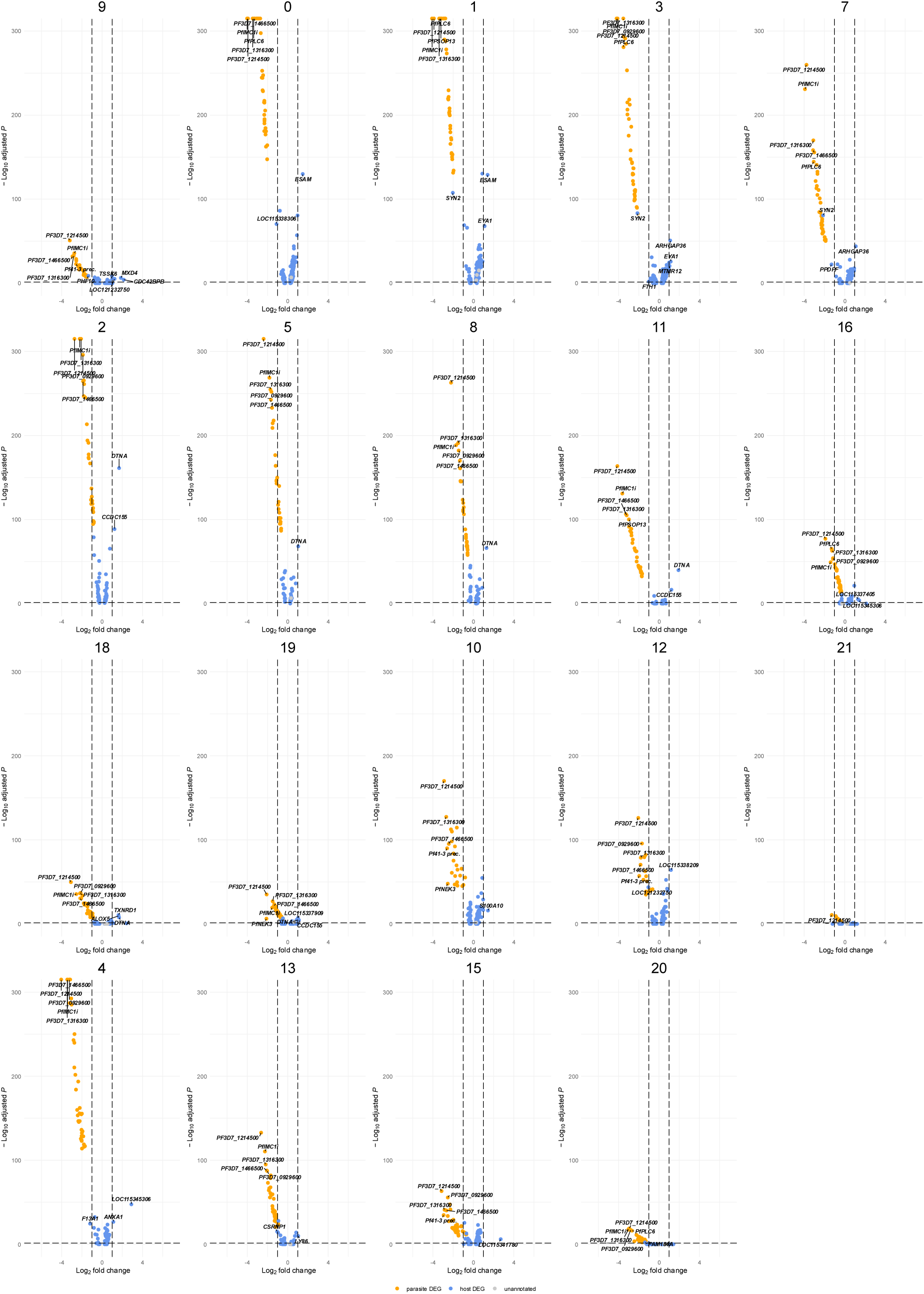
Differential gene expression across host clusters (top five host and parasite DEGs of LP versus HP are labelled). Dotted lines delimit cut-off-values (p-value (adjusted) ≤ 0.05; log₂-FC (averaged) ≥ 1). Negative and positive values for log₂-FC represent differential expression under HP versus LP respectively. Clusters are ordered by cell-type.

**Figure S6:**
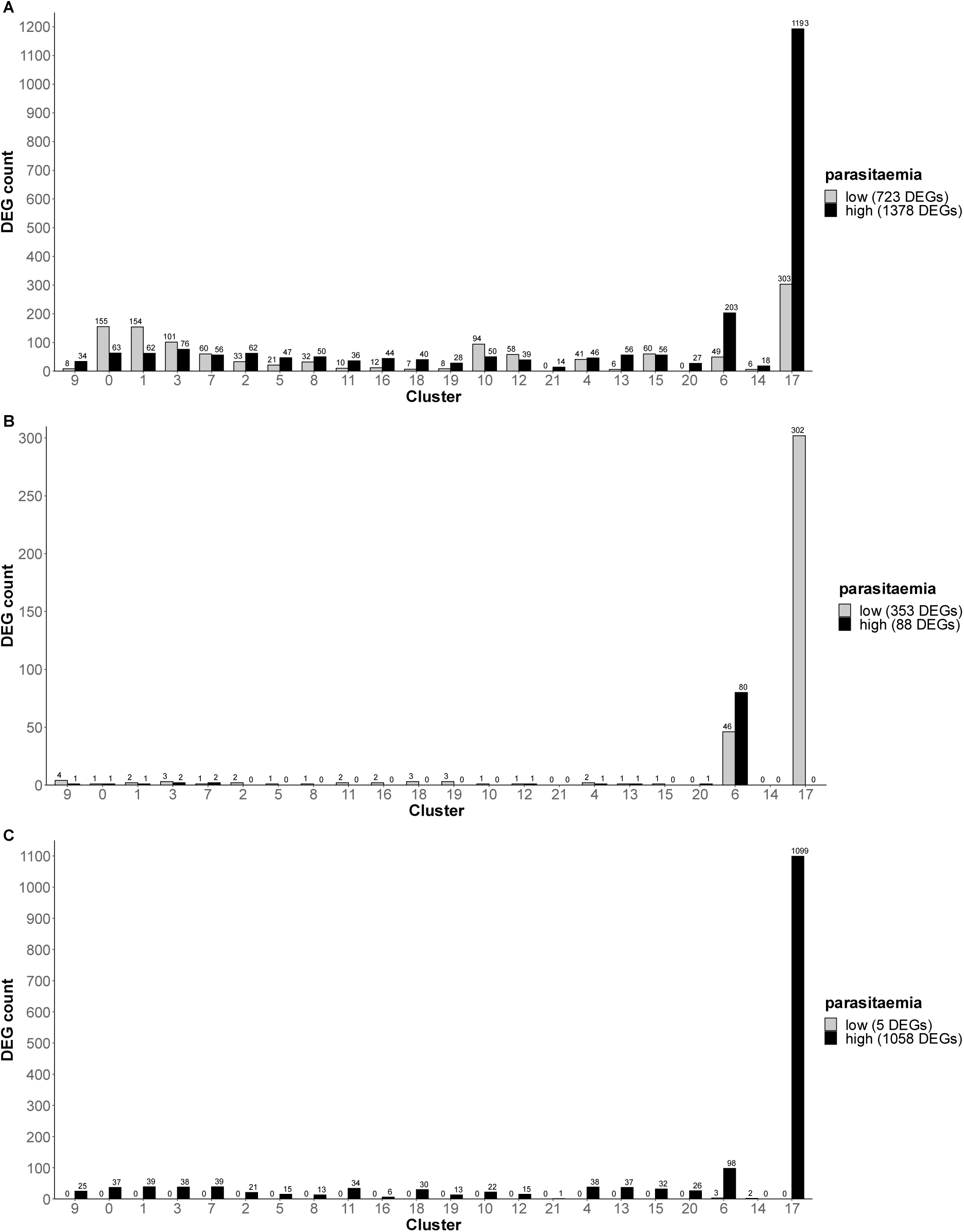
(A) Differentially expressed genes comparing LP versus HP per cluster (cut-off: p-value (adjusted) ≤ 0.05). (B) Strongly differentially expressed host genes (cut-off: p-value (adjusted) ≤ 0.05; log₂-FC (averaged) ≥ 1) comparing LP versus HP per cluster. Unannotated genes were omitted (C): Strongly differentially expressed parasite genes (cut-off: p-value (adjusted) ≤ 0.05; log₂-FC (averaged) ≥ 1) comparing LP versus HP per cluster. Unannotated genes were omitted. Clusters are ordered by cell type.

**Figure S7:**
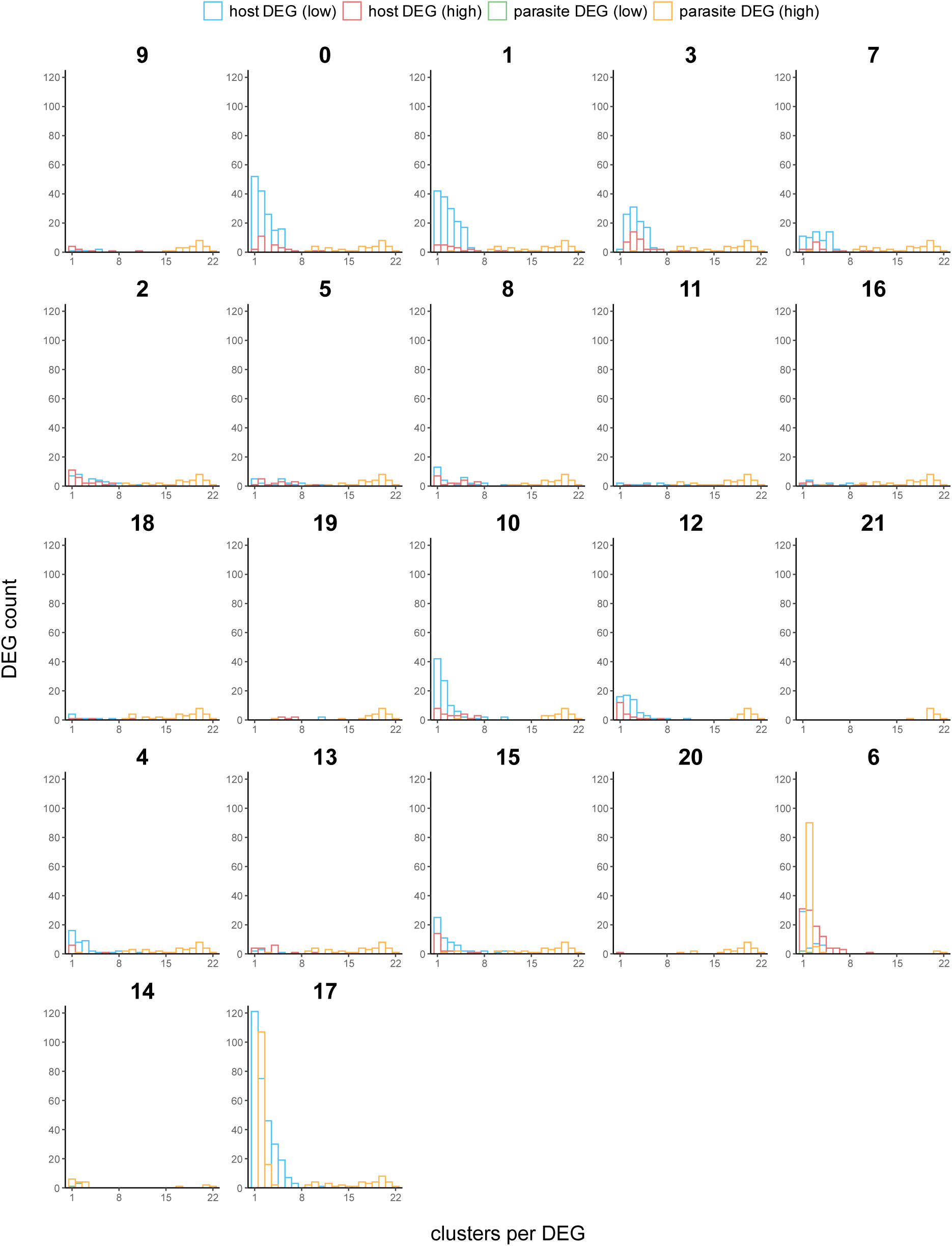
Redundancy of differentially expressed genes (cut-off: p-value (adjusted) ≤ 0.05). DEG counts are plotted against the number of clusters a gene is differentially expressed in. Unannotated genes were omitted.

